# Genome sequence of the medicinal plant *Tropaeolum majus* provides insights into flavonoid biosynthesis

**DOI:** 10.1101/2024.11.01.621629

**Authors:** Ronja Friedhoff, Ute Wittstock, Boas Pucker

## Abstract

*Tropaeolum majus* is a very popular species around the world with an enormous number of commercially available varieties displaying various flower color patterns and growth characteristics. It is rich in phytochemicals such as glucotropaeolin, hydroxycinnamic acid derivatives, and flavonol glycosides. Here, we report a highly continuous genome sequence of *T. majus* and a comprehensive annotation of protein encoding genes suitable for comparative genomics. The potential for the exploration of individual gene functions in this valuable plant is demonstrated by an analysis of the flavonoid biosynthesis genes and their transcriptional regulators. Important players of the flavonol biosynthesis, including structural genes and a transcription factor, were identified that are required to produce precursors of phytomedically relevant flavonol glycosides. The genome sequence does not reveal an ortholog of the leucoanthocyanidin reductase encoding gene (*LAR*), which aligns with previous reports about the absence of this gene in many Brassicales species.

## Introduction

The Tropaeolaceae family is native to South and Central America and represents one of the most ancient families of the Brassicales order. Its last common ancestor with the more recent core Brassicales has been dated to about 100 million years ago and its separation from Akaniaceae in the same clade to about 75 million years ago (Cardinal-McTeague *et al*., 2016). According to molecular-phylogenetic analyses, the about 90 species of the Tropaeolaceae can all be assigned to one genus, *Tropaeolum* (nasturtium), with several sections (Andersson × Andersson, 2000). Plants of this family are fleshy vines characterized by their large, mostly monosymmetric flowers with a nectar spur and their twining petioles (Missouri Botanical Garden, 2024). Because of their eye-catching flowers, several species, including *Tropaeolum majus, T. peregrinum* and *T. speciosum*, are grown as ornamental plants in gardens and parks. *T. tuberosum* (mashua) is grown in the Andes for its tuberous roots which are consumed by humans as a rich source of starch, protein, and fibers (Acurio *et al*., 2024).

*T. majus* is a very popular species around the world with an enormous number of commercially available varieties with various flower color patterns and growth characteristics. Its flowers are edible, have a spicy taste and are, for example, used to decorate salads. It is also used as a medicinal plant (Albrecht *et al*., 2023). For example, a preparation composed of *T. majus* herbal powder and *Armoracia rusticana* root powder has been approved in Germany for treatment of acute inflammatory diseases such as respiratory tract and urinary infections. The entire plant is rich in phytochemicals such as glucotropaeolin, hydroxycinnamic acid derivatives, and flavonol glycosides (Česlová *et al*., 2023). Apart from its horticultural and medicinal uses, *T. majus* has been subject of scientific research in various fields of plant biology. In the course of studies on cell wall biosynthesis, RNA-seq data have been generated from seeds and have been used to identify additional enzymes involved in xyloglucan biosynthesis (Jensen *et al*., 2012). However, a genome sequence, which would enable in-depth studies of primary and specialized metabolite biosynthesis pathways in *T. majus*, their regulation and evolution and further breeding efforts of this and other important species of the genus, is still missing. Here, we report a highly continuous genome sequence of *T. majus* and an annotation of the protein encoding genes. This genomic data set enabled a systematic investigation of the flavonoid biosynthesis in this valuable plant.

## Results & Discussion

### Genome sequence and annotation of *T. majus*

A total of 97.7 Gbp of ONT sequencing data was produced. Multiple assembly attempts were conducted (**Table 1**). Finally, the NextDenovo assembly was selected as the final genome sequence due to higher contiguity and a higher completeness. No bacterial or fungal contamination contigs were detected in this assembly. This is also supported by the GC content of 35% that is close to the average observed for other plant genome sequences (Pucker × Brockington, 2018). With an N50 of 35 Mbp, the continuity of this assembly exceeds the values reported for most previously released plant genome sequences (Marks *et al*., 2021) and provides a solid basis for comparative genomics. Synteny analyses and the discovery of biosynthetic gene clusters are facilitated with the release of this genome sequence.

**Table 1:** Statistics of the *T. majus* genome sequence assemblies.

|  | NextDenovo | Shasta |
| --- | --- | --- |
| Number of contigs | 149 | 168 |
| Total assembly size | 872.9 Mbp | 808.5 Mbp |
| Largest contig | 69.5 Mbp | 60.9 Mbp |
| N50 | 35 Mbp | 26.2 Mbp |
| BUSCO v3 | C:96.0%[S:92.0%,D:4.0%],<br>F:0.6%,M:3.4%,n:2326 | C:95.1%[S:91.1%,D:4.0%],<br>F:0.7%,M:4.2%,n:2326 |

Multiple gene prediction approaches were conducted based on the assembled genome sequence (**Table 2**). The annotation produced by GeMoMa based on structures of protein coding genes in other plant species performed best based on BUSCO results and was selected as the final annotation. The number of 32,924 predicted protein encoding genes is close to the average observed in plants (Pucker & Brockington, 2018). A prediction of the gene functions was generated based on sequence similarity to *Arabidopsis thaliana* sequences and information available from TAIR10/Araport11 (Pucker *et al*., 2024).

**Table 2:**
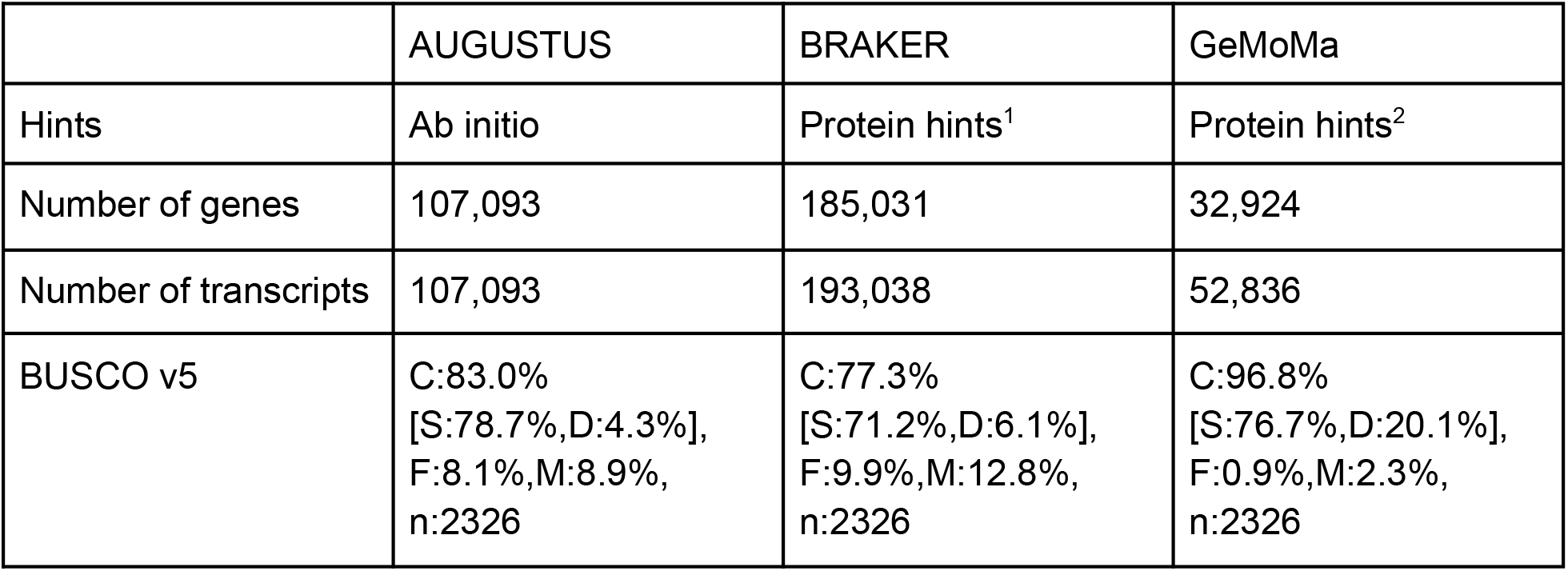
Selected gene prediction results for the *T. majus* genome sequence. Different protein hints were used for: ^1^ BRAKER (Viridiplantae) and ^2^ GeMoMa (*Carica papaya, Theobroma cacao, Arabidopsis halleri, A. thaliana, Eutrema salsugineum*, and *Thlaspi arvense*).

|  | AUGUSTUS | BRAKER | GeMoMa |
| --- | --- | --- | --- |
| Hints | Ab initio | Protein hints<sup>1</sup> | Protein hints<sup>2</sup> |
| Number of genes | 107,093 | 185,031 | 32,924 |
| Number of transcripts | 107,093 | 193,038 | 52,836 |
| BUSCO v5 | C:83.0%<br>[S:78.7%,D:4.3%],<br>F:8.1%,M:8.9%,<br>n:2326 | C:77.3%<br>[S:71.2%,D:6.1%],<br>F:9.9%,M:12.8%,<br>n:2326 | C:96.8%<br>[S:76.7%,D:20.1%],<br>F:0.9%,M:2.3%,<br>n:2326 |

### Investigation of the flavonoid biosynthesis in *T. majus*

Genes for all steps of the core flavonoid biosynthesis were identified in the *T. majus* gene set (**Table 3, Fig. 1**). The presence of flavonoid biosynthesis genes in *T. majus* is expected given previous reports about the presence of flavonoids (Garzón *et al*., 2015; Česlová *et al*., 2023). Especially relevant for the biosynthesis of previously reported flavonol glycosides in *T. majus* (Garzón *et al*., 2015; Česlová *et al*., 2023) are the genes *CHS, CHI, F3H, F3’H, F3’5’H*, and *FLS* (Winkel-Shirley, 2001; Pucker *et al*., 2020). Although not all land plants harbor *F3’5’H*, the detection in *T. majus* aligns with reports of myricetin derivatives (Garzón *et al*., 2015) which requires hydroxylation reactions that can be catalyzed by F3’5’H. A gene encoding a member of the well known flavonol biosynthesis regulating R2R3-MYB subgroup 7 (Stracke *et al*., 2007, 2010) was identified which could be a target for future attempts to engineer the flavonol biosynthesis in this medicinal plant.

**Table 3:**
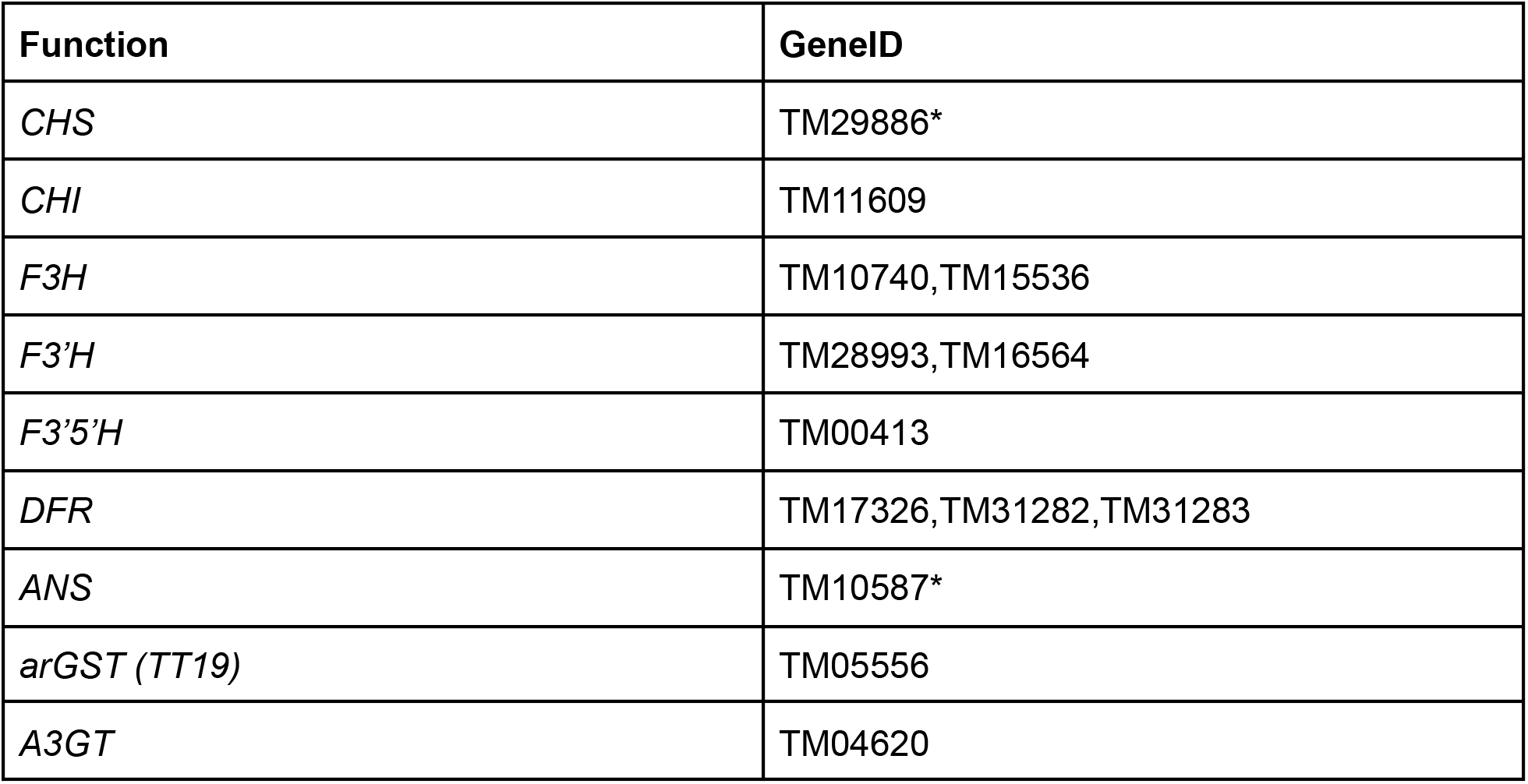
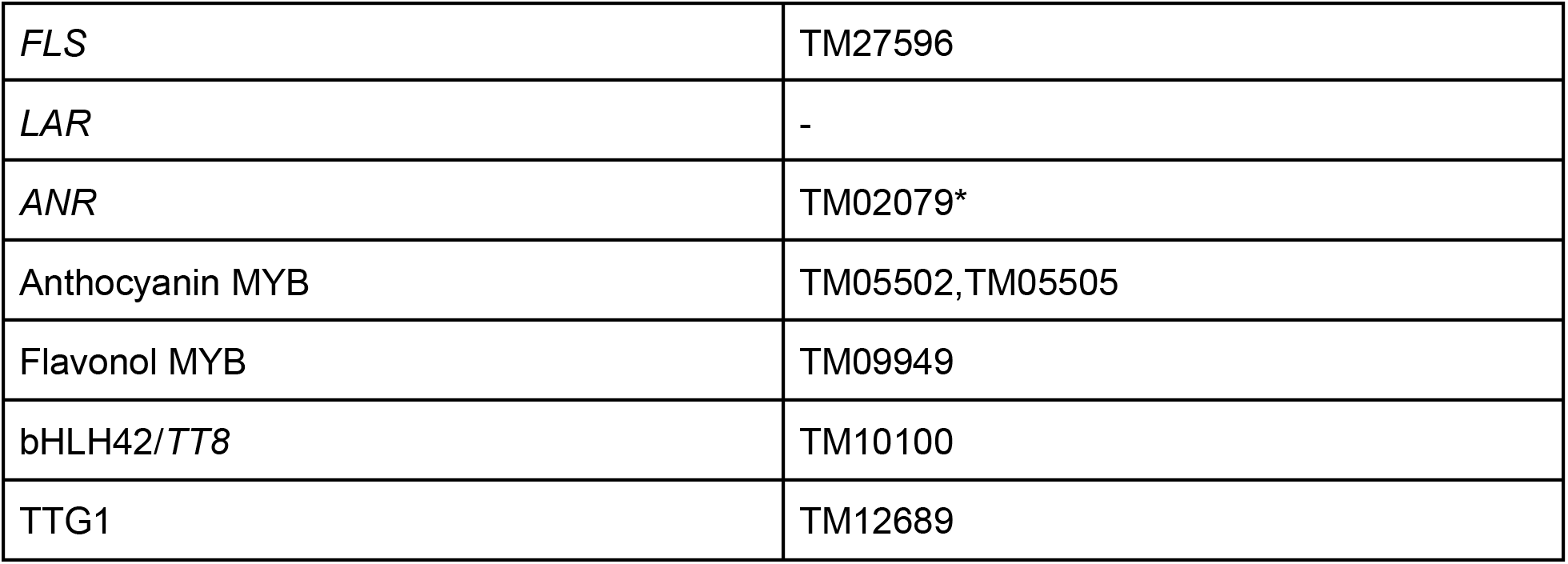
Annotation of the flavonoid biosynthesis genes in *T. majus*. Candidates lacking a residue in their predicted amino acid sequence that is considered crucial for their predicted function are marked with an asterisk.

**Fig. 1:**
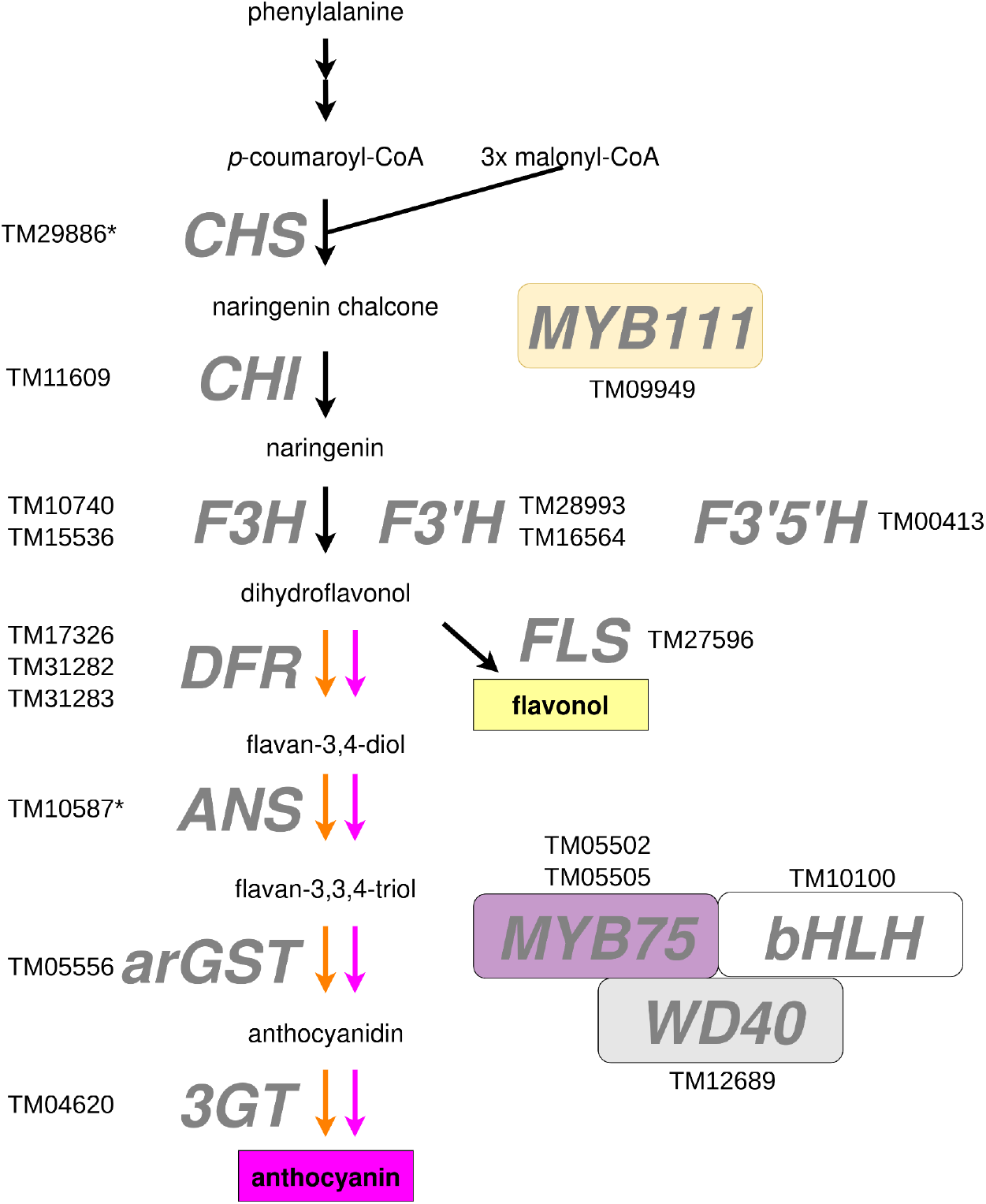
Identified candidate genes in the flavonoid biosynthesis. Candidates lacking an amino acid residue in the derived amino acid sequence that is considered crucial for their predicted function are marked with an asterisk. The layout of this figure is based on (Horz *et al*., 2024). Activity of the flavonoid biosynthesis genes was assessed based on RNA-seq data (SRR25909845) belonging to seed samples (**Fig. 2**). Many genes of the anthocyanin biosynthesis are also part of the proanthocyanidin biosynthesis, which could be active in seeds of *T. majus*. Previous studies have reported important roles and substantial levels of proanthocyanidins in seeds of *A. thaliana* and *Vitis vinifera* (Nakamura *et al*., 2003; Debeaujon *et al*., 2003). It is also plausible that no flavonols are produced in the seeds which could explain the inactivity of *FLS* and its regulator.

**Fig. 2:**
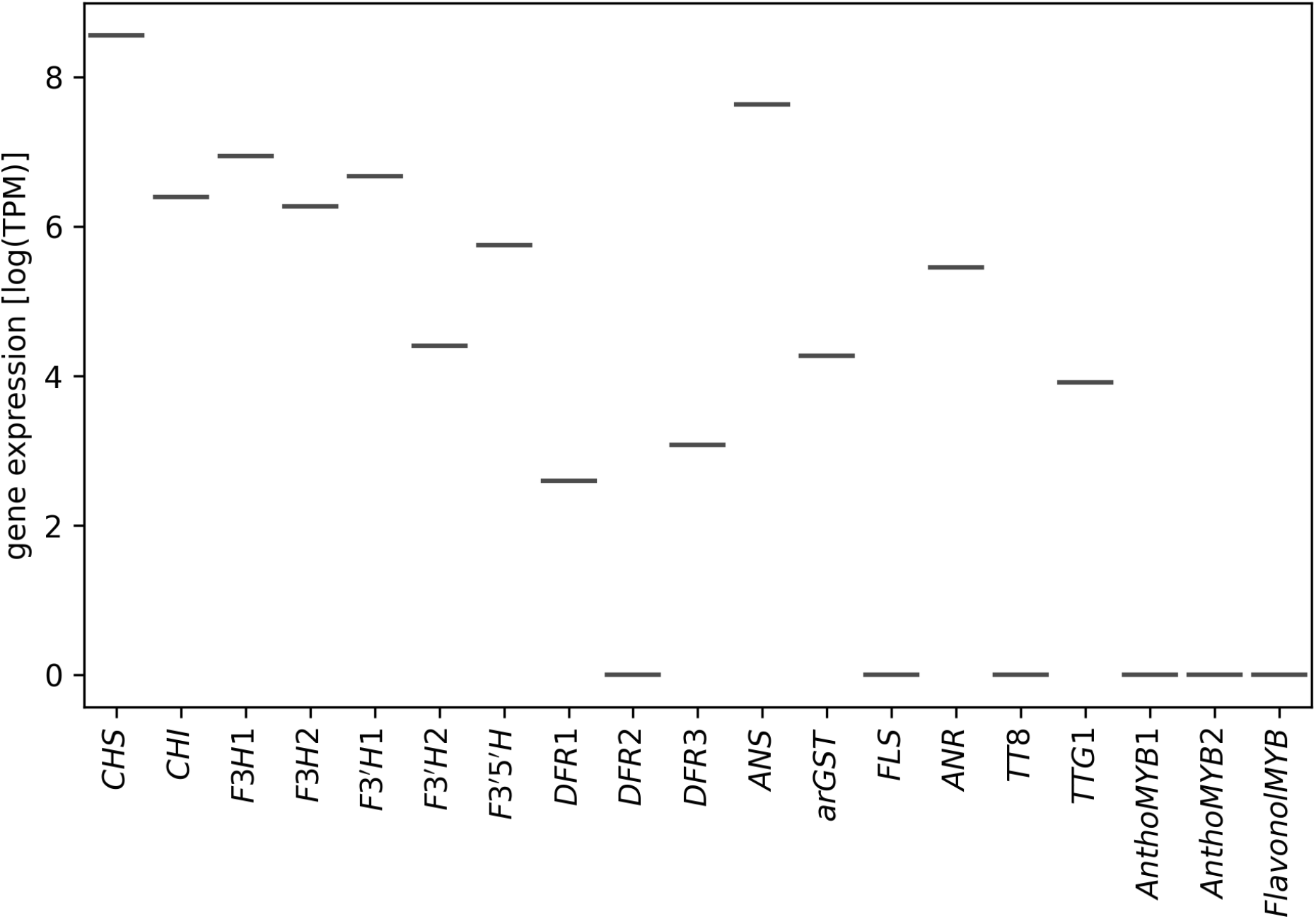
Expression of flavonoid biosynthesis associated genes in seeds of *T. majus* (SRR25909845). Transcripts per million (TPMs) are displayed as log-transformed values.

No gene encoding leucoanthocyanidin reductase (LAR) was detected in this *T. majus* gene set. Given the phylogenetic position of *T. majus* within the Brassicales, this aligns with previous reports about the absence of LAR in the Brassicales (Nesi *et al*., 2009; Pucker *et al*., 2020; Lu, 2024). The functional and ecological consequences of the absence of LAR remain unknown. Although *LAR* is considered an important gene in the proanthocyanidin biosynthesis (Tanner *et al*., 2003; Pang *et al*., 2007; Liu *et al*., 2016), many Brassicales species are well known for their ability to accumulate oligomeric proanthocyanidins. For example, the proanthocyanidin biosynthesis in *A. thaliana* was elucidated through a collection of transparent testa mutants that harbor a block in one of the proanthocyanin biosynthesis genes (Appelhagen *et al*., 2014).

## Materials & methods

### DNA extraction

*T. majus* L. (Accession TRO1, ID 47139) fruits were obtained from Genebank Gatersleben of the Leibniz Institute of Plant Genetics and Crop Plant Research, Germany. After soaking in water for 2 h, fruits were placed on Vermiculite and left in a controlled environment chamber at 22°C, 65 % humidity with a 10 h photoperiod (120 µmol m^-2^ s^-1^ photosynthetically active radiation) for a week. Seedlings were then transferred to potting substrate (germination potting compost (Compo, Münster, Germany) supplemented with 10 % (v/v) sand, 10 % (v/v) Perligran (Knauf Perlite, Germany), 2 g/l Triabon (Compo) and 1 g/l Sierrablen (Scotts, Heerlen, NL)) which had been steamed at 100°C for two hours and treated with Neudomück^®^ (W. Neudorf GmbH KG, Emmerthal, Germany). Seventeen days after sawing, the photoperiod was changed to 16 h light. Plantlets were transferred to larger pots filled with steamed and Neudomück^®^-treated potting compost with 10 % (v/v) Perligran on day 32 after sawing The largest leaves with parts of the petioles were harvested from a single plant at two time points (35 and 40 days after sawing), placed on ice and used immediately to isolate genomic DNA.

Leaf material was homogenized by grinding in liquid nitrogen. High molecular weight DNA was extracted using a modified CTAB-based protocol as previously described (Siadjeu *et al*., 2020). DNA quality was assessed via NanoDrop measurement, agarose gel electrophoresis, and Qubit measurement as previously described (Horz *et al*., 2024). Short DNA fragments were depleted with the Short Read Eliminator kit (Pacific Biosciences) following the suppliers instructions.

### ONT sequencing

Nanopore sequencing libraries were prepared with the SQK-LSK109 ligation-based kit (Oxford Nanopore Technologies) using 1 µg of DNA per library and following the suppliers’ instructions. ONT sequencing was performed on a MinION-Mk1B with R9.4.1 flow cells. Flow cell washing was performed upon blockage of a large proportion of nanopores followed by loading of a fresh library to achieve optimal performance of the flow cells. Basecalling of the raw sequencing data was performed with Guppy v6.4.6 (ONT) in high accuracy on a GPU in the de.NBI cloud. The customized Python script FASTQ_stats.py was applied to calculate statistics of the obtained sequencing data (https://github.com/bpucker/GenomeAssembly).

### Genome sequence assembly

Initial assemblies were generated with Shasta v0.11.1 (Shafin *et al*., 2020) to assess coverage of the sequencing data. The final assembly was generated based on all reads and NextDenovo v2.5.2 (Hu *et al*., 2024) with the parameters genome_size = 800m, read_cutoff = 10k, seed_depth=45. The assembly statistics were compared with contig_stats3.py (Meckoni *et al*., 2023). BUSCO v3.0.2 (Simão *et al*., 2015; Manni *et al*., 2021) was run in the genome mode to assess the completeness of both assemblies. The assembly generated by NextDenovo was selected as the representative genome sequence of *Tropaeolum majus*. The assembly was processed with clean_genomie_FASTA.py to reduce the sequence names to short identifiers without any special characters. Next, the genome sequence was screened for bacterial/fungal contamination contigs using assembly_wb_screen.py (Meckoni *et al*., 2023).

### Structural and functional annotation

Different approaches were conducted to predict protein encoding genes based on the assembled genome sequence. All results were compared with BUSCO v3.0.2 (Simão *et al*., 2015; Manni *et al*., 2021) run in the protein mode with embryophyta_odb10 as reference. RNA-seq reads of *T. majus* (SRR25909845) were retrieved from the Sequence Read Archive (Katz *et al*., 2022) and aligned to the assembled genome sequence with STAR v2.7.11b (Dobin *et al*., 2013; Dobin & Gingeras, 2015). BRAKER v3 (Gabriel *et al*., 2023) was run with the aligned RNA-seq reads as hints. Another BRAKER run was supplied with protein hints derived from Viridiplantae (NCBI, 2023-05-01). AUGUSTUS v3.3 (Stanke *et al*., 2006) was run in *ab initio* mode using previously optimized parameters for the detection of non-canonical splice sites (Pucker *et al*., 2017). GeMoMa v1.9 (Keilwagen *et al*., 2016, 2019) was run with protein hints derived from the following species *Carica papaya* (GCF_000150535.2) (Ming *et al*., 2008), *Theobroma cacao* (GCF_000208745.1) (Argout *et al*., 2011), *Arabidopsis halleri* (GCA_964271285.1), *A. thaliana* (GCF_000001735.4) (The Arabidopsis Genome Initiative, 2000; Sloan *et al*., 2018), *Eutrema salsugineum* (GCF_000478725.1) (Yang *et al*., 2013), and *Thlaspi arvense* (GCA_911865555.2) (Nunn *et al*., 2022). The structural annotation produced by GeMoMa based on the comprehensive set of protein hints was selected as representative annotation. A corresponding functional annotation was generated with construct_anno.py (Pucker & Iorizzo, 2023) based on knowledge about gene functions in *A. thaliana* (Lamesch *et al*., 2012; Cheng *et al*., 2017).

### Gene expression analysis

Kallisto v044 (Bray *et al*., 2016) was used to process the publicly available RNA-seq data. Customized Python scripts were applied for the generation of a final count table (Pucker × Iorizzo, 2023). A figure summarizing the expression of flavonoid biosynthesis genes was generated based on normalized expression data (transcripts per million, TPMs) with a Python script using the matplotlib, pandas, scipy, numpy, and seaborn modules (Hunter, 2007; Virtanen *et al*., 2020; Harris *et al*., 2020; Waskom, 2021; Pucker & Iorizzo, 2023; The pandas development team, 2024).

## Data availability

All data sets underlying this study are publicly available. Sequencing data have been deposited at the European Nucleotide Archive (PRJEB68329). The assembled genome sequence and corresponding annotation have been published via LeoPARD (https://leopard.tu-braunschweig.de/receive/dbbs_mods_00078092).

## Acknowledgements

This work was supported by the BMBF-funded de.NBI Cloud within the German Network for Bioinformatics Infrastructure (de.NBI) (031A532B, 031A533A, 031A533B, 031A534A, 031A535A, 031A537A, 031A537B, 031A537C, 031A537D, 031A538A). We thank GeneBank at the IPK Gatersleben for providing *T. majus* fruits, Anita Backenköhler (Institute of Pharmaceutical Biology) for raising plants and all members of the research group Plant Biotechnology and Bioinformatics for discussion and support.

## Author contributions

BP and UW conceived the project. RF extracted DNA and performed the sequencing. RF and BP conducted the bioinformatic analyses. UW and BP wrote the manuscript. RF, UW, and BP reviewed the final version of the manuscript and consented to its submission.

